# Genome-wide shifts in histone modifications at early stage of rice infection with Meloidogyne graminicola

**DOI:** 10.1101/2020.07.06.190538

**Authors:** Mohammad Reza Atighi, Bruno Verstraeten, Tim De Meyer, Tina Kyndt

## Abstract

Epigenetic processes play a crucial role in the regulation of plant stress responses, but their role in plant-pathogen interactions remains poorly understood. Although histone modifying enzymes have been observed to be deregulated in galls induced by root-knot nematodes (RKN, *Meloidogyne graminicola*) in rice, their influence on plant defence and their genome-wide impact have not been comprehensively investigated.

In this study, we applied 3 chemical inhibitors of histone modifying enzymes on rice 24h before inoculation with RKN. Despite their distinct described effects on histone modifications, application of different concentrations led in all cases to reduced susceptibility to RKN. Infection assays on two overexpression lines of histone lysine methyltransferases and one histone deacetylase showed contrasting results in susceptibility, indicating that each histone mark has a specific role in plant defence against RKN in rice. At genome-wide level, three histone marks, H3K9ac, H3K9me2 and H3K27me3 were studied by chromatin-immunoprecipitation (ChIP)-sequencing on RKN-induced galls at 3 days post inoculation. While levels of H3K9ac and H3K27me3 were strongly enriched, H3K9me2 was generally depleted in galls versus control root tips. Differential histone peaks were generally associated with plant defence related genes. In conclusion, our results indicate that histone modifications respond dynamically to RKN infection, and specifically target plant-defence related genes.

**One-sentence summary:** Post-translational histone modifications show a dynamic genome-wide response to root-knot nematode infection in rice and are specifically associated with plant defence genes.

## Introduction

In plants, as in other eukaryotes, DNA is wrapped around histone proteins and the resulting DNA-histone complex or nucleosome is the basic repeating unit of chromatin. The histone tails protruding from the nucleosome core can be modified by addition of chemical groups, mainly acetyl and methyl groups, affecting the physical accessibility of DNA to the transcriptional machinery of the cell (Berger, 2007; Lawrence et al., 2016). Fine-tuning of gene expression is obtained by an interplay between different types and levels of post-translational histone modifications (Schones and Zhao, 2008; Zhou et al., 2011). Some histone marks are associated with transcriptional activation, e.g. acetylation of lysine residues on histone H3 is correlated with gene activation and in some cases DNA repair. In contrast, depending on the residue, H3 methylation can be associated with transcriptional activation (lysine 4 and 36) or repression (lysine 9 and 27) (Fuchs et al., 2006; Berger, 2007; Armstrong, 2014). Both lysine and arginine can be methylated and up to three methyl groups can bind to each residue. The histone acetylation level is balanced by the activity of histone acetyltransferases (HATs) and histone deacetylases (HDACs or HDAs). The relatively small genome of rice contains 8 HATs and 19 HDACs (Pandey et al., 2002; Liu et al., 2012; Zhou et al., 2013). The rice genome also contains 37 Su(var)3-9/Enhancer of Zeste/Trithorax (SET)-domain containing histone methyltransferases also named the SET domain-containing group (SDG), eight protein arginine methyltransferases (PRMTs) and 24 Jumonji C domain-containing histone-lysine demethylases (JmjC-KDMs) which regulate the histone methylome (Zhou et al., 2013).

The genome-wide distribution of histone marks has been mainly investigated in unstressed plants. Du et al. (2013) found H3K9ac to be mainly present (73.4%) in genic regions [promoter, 5’ untranslated region (UTR), 3’ UTR, exons and introns] of the rice genome. They also observed that H3K9ac is associated with 781 transposable-element genes (TE genes) and 19,616 non-TE genes (Du et al., 2013). In another study, the distribution patterns of three histone marks, H3K4me3, H3K9ac and H3K27me3 showed overall enrichment in genic and euchromatic regions. Further, they observed that out of 41,043 non-TE genes in the TIGR database, 25,207 (61.4%), 26,623 (64.9%), and 17,211 (41.9%) were associated with H3K4me3, H3K9ac, and H3K27me3, respectively. In contrast, these histone marks were associated with less than 5.2% of TE–related genes in the same database (He et al., 2010).

Modifications of histone tails are a major mechanism in gene expression regulation, and their role in plant responses to environmental stimuli and pathogen infection has been described (Zhou et al., 2005; Kim et al., 2008; Jaskiewicz et al., 2011; Choi et al., 2012; Li et al., 2013; Dutta et al., 2017; Vijayapalani et al., 2018). For example, upon infection of *Paulownia fortunei* with phytoplasma, 1788 and 939 genomic regions were hyper- and hypoacetylated, respectively for H3K9ac (Yan et al., 2019). Application of salicylic acid (SA)- analogue Benzothiadiazole (BTH) induced H3K9ac enrichment in the promoter of *pathogenesis-related gene 1* (*PR1*) (Lopez et al., 2011). In the same line, pathogen infection or treatment with agents that induce plant resistance tends to lead to higher expression levels of HATs, as was for example observed upon aplication of the hormones abscisic acid (ABA) and SA (Liu et al., 2012). Similarly, Singh et al., 2014 showed that the Arabidopsis *hat-1* mutant plants failed to show enhanced resistance against *Pseudomonas syringae* pv *tomato* DC3000 upon priming by recurrent heat, cold, or salt stress compared to wild type plants which could retain the memory of stress (Singh et al., 2014).

On the other hand, expression of HDA19 - the best-studied HDAC in Arabidopsis - is induced by wounding, infection by fungal pathogen *Alternaria brassicicola*, and application of jasmonic acid (JA) and ethylene (ET) (Zhou et al., 2005). Overexpression of *HDA19* confers higher resistance in plants against *A. brassicicola*, whereas the *hda19-*deficient mutant shows increased susceptibility to this pathogen through differential regulation of genes involved in the JA and ET pathways (Zhou et al., 2005). Similarly, Kim et al., 2008 showed that overexpression of *HDA19* results in enhanced resistance against *P. syringae*, while the deficient mutant was more susceptible to infection, through compromised activation of WRKY38 and WRKY62, which are negative regulators of plant defence (Kim et al., 2008). However, in another study it was demonstrated that HDA19 represses the SA-mediated defence response in *Arabidopsis*, and the *hda19*-mutant showed enhanced resistance against *Pseudomonas syringae* pv *tomato* DC3000 (*Pst* DC3000), correlated with enhanced expression of *PR* genes (Choi et al., 2012). Similarly, the *hda6* mutant showed increased levels of acetylation which led to enhanced resistance to *Pst* DC3000 by increased activation of *PR1, PR2* and *PR4* (Wang et al., 2017). Given its clear role in plant defence, it is not unexpected that pathogens actively interfere with histone acetylation. An effector of the sugar beet cyst nematode *Heterodera schachtii* alters acetylation levels in *Arabidopsis* by inhibiting HDACs, leading to acetylation of rRNA genes and their activation (Vijayapalani et al., 2018).

Next to histone acetylation, also other marks are involved in plant defence. In bean plants infected with rust causing pathogen *Uromyces appendiculatus*, next to H4K12ac also H3K9me2 marks were associated with several important plant defence genes, including WRKY, bZIP, MYB transcription factors, chitinase, calmodulin and leucine-rich repeat (LRR) genes. At the genome-wide level, it was demonstrated that peaks for both histone marks were mainly located in intergenic regions (Ayyappan et al., 2015). Similarly, JmjC DOMAIN-CONTAINING PROTEIN 27 (JMJ27), a H3K9me1/2 demethylase, was shown to be a positive regulator of plant defence against *Pst* DC3000 in Arabidopsis. JMJ27 negatively regulates WRKY25 (a repressor of defence) and positively regulates *PR* genes (Dutta et al., 2017). JMJ705 overexpression in rice confers enhanced resistance against *Xanthomonas oryzae* pv. *oryzae* by reduction in H3K27me2/3 levels, correlated with higher expression of defence related genes encoding for example peroxidases, JA-related genes and PR-proteins (Li et al., 2013).

The use of chemical inhibitors of histone modifying enzymes to counteract epigenetic malfunctions in human oncogenes has been studied extensively in cancer research (Heerboth et al., 2014). Recent studies have also shown the potential of chemical reagents in influencing epigenetic mechanisms in plants (Tanaka et al., 2008; Bond et al., 2009; Zhang et al., 2012; Zhang et al., 2013; Miwa et al., 2017). As HDAC-inhibitor, nicotinamide application induces the expression of *VERNALIZATION INSENSITIVE 3* (*VIN3*) in *Arabidopsis*, causing flowering and repression of flowering locus C (Bond et al., 2009). Nicotinamide also impacts defence, as its derivative, nicotinamide mononucleotide, accumulates in barley cultivars that are resistant to *Fusarium*. Pre-treatment of plants with nicotinamide mononucleotide confers enhanced resistance to *Fusarium graminearum* in *Arabidopsis* leaves and flowers as well as in barley spikes (Miwa et al., 2017). Sulfamethazine (SMZ) belongs to the family of antibacterial sulfonamides that impair folate synthesis and thereby methyl supply in plants. Application of sulfamethazine on Arabidopsis reduces the level of H3K9me2 methylation and DNA methylation and consequently derepresses epigenetic silencing (Zhang et al., 2012). Fumarate inhibits histone demethylation at sites of DNA damage (Lees Miller, 2015). It induces resistance against *Pst* DC3000 in *Arabidopsis* without any direct antimicrobial effect (Balmer et al., 2018).

In this study, we aimed at assessing the impact of histone modifications on rice plant susceptibility to nematodes. Rice is the staple food of half the world’s population. With a relatively small and fully sequenced genome, it is an excellent model system for genomic studies in monocotyledonous plants. The root knot nematode *Meloidogyne graminicola* (*Mg*), is one of the most damaging nematodes attacking rice and other monocots (Bridge et al., 2005). After penetration of second stage juveniles of *Mg* into the rice roots, they establish a feeding site which consists of giant cells. Giant cell establishment causes root deformation, leading to symptoms called galls, which in the case of *Mg* are typically located at the root tip and are macroscopically visible from about 3 dpi (Mantelin et al., 2017).

In previous transcriptome studies conducted in our lab, expression of many histone modification enzymes was significantly altered in galls and giant cells induced by *Mg* in comparison with control tissues/cells (Kyndt et al., 2012; Ji et al., 2013). Here, we studied how histone modifications are affected in galls at very early time points after infection and what is their role in plant defence against *Mg*. First, we used chemical inhibitors and transgenic rice lines to study how histone acetylation and methylation influence rice susceptibility to nematode infection. Activation/over-expression lines of OsSDG740, OsSDG729 and OsHDA712 were selected for infection assays because a diverging modulation of their expression upon nematode infection was observed (Kyndt et al., 2012; Ji et al., 2013), suggesting their possible involvement in rice-*Mg* interactions. Second, using ChIP-Seq we studied changes in patterns of three histone marks, one activation (H3K9ac) and two suppression (H3K9me2 and H3K27me3) marks, to study which genomic regions are marked by activation and suppression marks upon nematode infection.

## Results

Upon root-knot nematode *Meloidogyne graminicola* infection in rice, a strong transcriptional deregulation occurs in many histone modifying genes at the early stage of infection, 3 dpi (Kyndt et al., 2012). At this moment, root-knot nematodes are initiating a feeding site at the rice root tips, which is correlated with strong effects on plant defence (Mantelin et al., 2017; Kyndt et al., 2012). About 80 genes encoding enzymes involved in histone lysine acetylation and methylation are differentially expressed upon nematode infection (Supplementary figure 1), indicating changes at the epigenomic level. In the current study, chemical inhibitors of histone modifying enzymes and transgenic lines over-expressing three histone modifying enzymes, with diverging patterns of expression upon inoculation (see arrows in Supplementary figure 1), were studied to evaluate if changes in the histone mark profile in rice would affect susceptibility to nematode infection. The genome-wide histone modification profile of three histone marks namely H3K9ac, H3K9me2 and H3K27me3 were subsequently analyzed in galls in comparison with uninfected root tips.

### Disruption of histone modification patterns alters susceptibility of rice plants to nematode infection

To investigate whether interference with histone modifications affects the response of rice plants to nematode infection, three chemical inhibitors namely nicotinamide (HDAC inhibitor), sulfamethazine (histone and DNA methyltransferase inhibitor) and fumaric acid (histone demethylase inhibitor) were used. The activity of these chemical inhibitors and their effect on plant development and defence was described in previous studies, using the same concentrations as applied here (Tanaka et al., 2008; Bond et al., 2009; Zhang et al., 2012; Zhang et al., 2013; Miwa et al., 2017). All chemicals were sprayed (Latzel et al., 2016; Miwa et al., 2017; Puy et al., 2018; Dawood et al., 2019) on plants 24 h before nematode inoculation. To rule out any direct effects on the nematodes, we executed foliar application of these chemicals including a surfactant to allow adequate uptake and systemic effects in the plants (Nahar et al., 2011). At 14 days after spraying, plants were harvested and number of galls and nematodes per plants were counted. Data were also expressed per milligram of dry root, to correct for any potential growth retardation effects induced by these chemicals in treated plants.

Two different concentrations of fumaric acid, 2.5 mM and 5 mM, were applied, showing no observable developmental or growth retardation effects in plants (Figure 1A). However, the numbers of galls and nematodes decreased significantly, and in a concentration-dependent manner, whether expressed per plant or per milligram of root (Figure 1B and C). Sulfamethazine application caused negative effects on plant growth at 2000 µM but not at 200 µM (Figure 1D). Number of galls and nematodes were significantly decreased, whether expressed per plant (Figure 1D) or per milligram of root (Figure 1E, F). We applied two concentrations of nicotinamide, 1 mM and 10 mM. The lower concentration slightly promoted root development, as previously observed in wheat roots and shoots (Youssef et al., 1989). However, this application had no significant effect on number of galls or nematodes compared to untreated control plants (Figure 1G, H and I). The 10 mM concentration caused a significant retardation in root and shoot development compared to the 1 mM treated and control plants (Figure 1G). The number of nematodes was significantly lower in treated plants, per plant and per milligram of root, but no significant differences were observed for number of galls (Figure 1 H and I).

**Figure 1.**
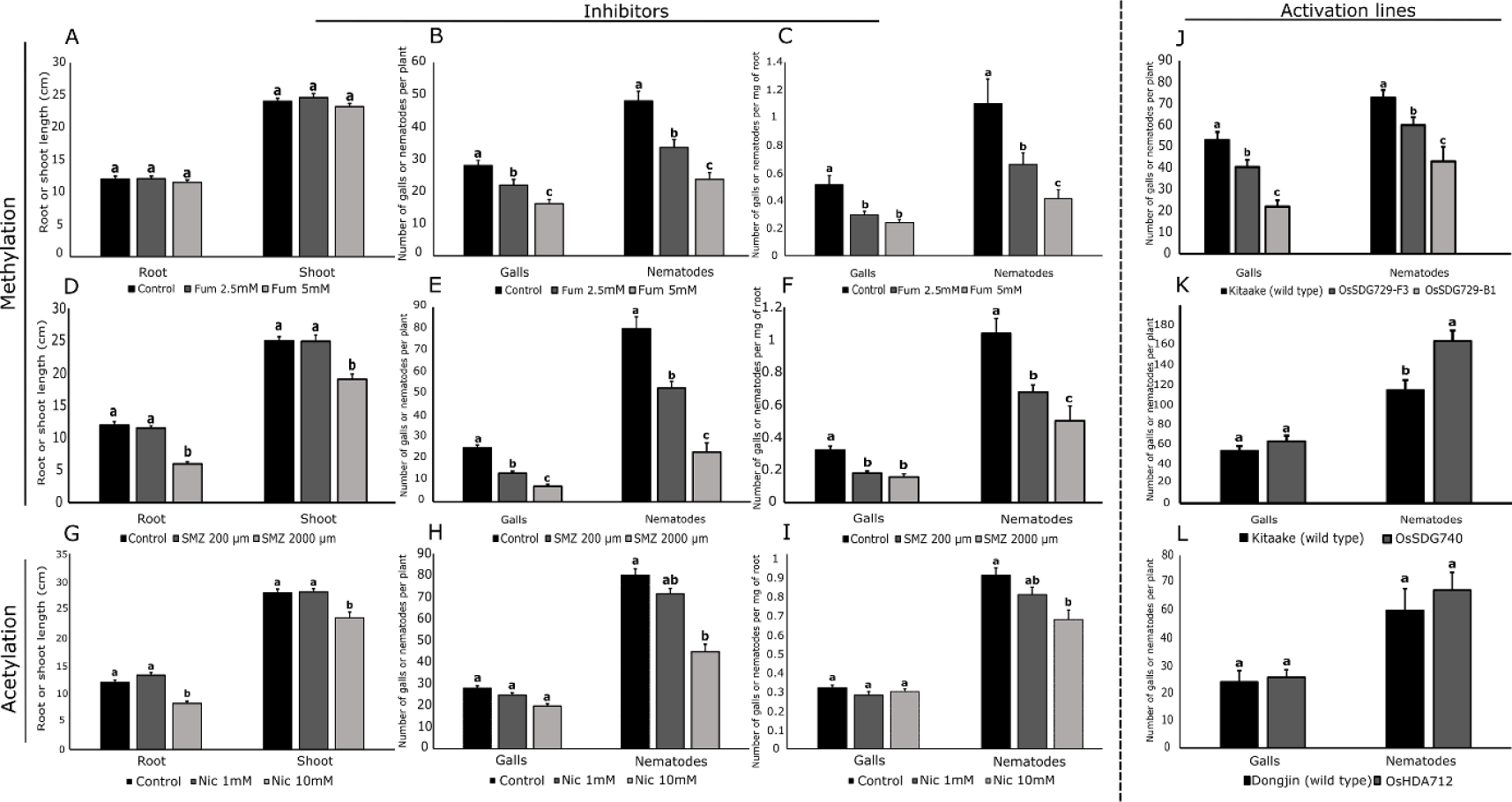
Effect of chemical or genetic interference with histone modifications on rice growth and susceptibility for root-knot nematode infection. Plants were treated with different chemicals at 24 h before inoculation with 300 J2 of *M. graminicola* per plant. Root and shoot length as well as number of galls were evaluated 14 days later. (**A - C**) Effect of fumaric acid on rice growth and susceptibility to nematodes, (**E - F**) Effect of sulfamethazine on plant growth and susceptibility to nematodes, (**G - I**) Effect of nicotinamide on plant growth and susceptibility to nematodes (n = 20), (**J, K and L**) Effect of overexpression of *OsSDG729, OsSDG740* and *OsHDA712* on susceptibility to nematodes (n = 18). For all subpanels different letters denote significant differences (p <0.05). Error bars indicate SEM.

To further study the effect of histone methylation/acetylation on plant defence against nematodes, we used two lines over-expressing histone lysine methyltransferases of the SDG family (OsSDG729, OsSDG740) and 1 HDAC activation line (OsHDA712). These genes show diverging expression profiles upon RKN infection (supplementary figure 1; black arrows). While OsHDA712 is strongly activated, OsSDG729 shows only minor differences in expression in galls versus control root tips, while OsSDG740 is repressed in galls. Despite the 11.5-128 fold induction of the target genes, no significant developmental defects were observed for any of these lines (Supplementary Figure 2). However, infection assays revealed that overexpression of OsSDG729 confers significantly reduced susceptibility to nematode infection, at about 23.9%-58.4% and 18.1%-40.9% for number of galls and nematodes, receptively compared to wild-type plants (Figure 1J). On the other hand, the OsSDG740 activation line shows enhanced levels of nematode penetration, with the number of nematodes being significantly increased by 43.1%, while the number of galls was not significantly changed (only 17.6%) in comparison with wild type plants (Figure 1K). The OsHDA712 activation line on the other hand showed no difference in susceptibility to root-knot nematodes compared to wild type plants (Figure 1L). These data indicate that interfering with plant histone marks may be associated with changes in plant defence against *Mg* in rice. They also highlight that each of these genes potentially control a subset of downstream genes which leads to different effects on plant susceptibility.

### Nematode–induced galls feature increased levels of H3K9ac

The involvement of H3K9 acetylation (H3K9ac), a gene activation mark, in plant defence against fungi and bacteria was studied before (Yan et al., 2019). Here, galls at 3 dpi and corresponding root tips were collected and were used for chromatin immunoprecipitation following DNA sequencing. In total 1062 peak regions were significantly differentially modified in galls compared to control root tips (Supplementary file 3). We observed H3K9 hyperacetylation in galls compared to roots tips (Figure 2A). The peak distribution showed an increase of this histone mark around the TSS of protein coding genes. However, right at the TSS it is less pronounced compared to immediate adjacent regions, similar to observations in human samples (Klein et al., 2019) (Figure 2B). To see whether these histone peaks are associated with genomic elements (e.g. promoter, gene bodies and TEs), their association was evaluated. H3K9ac peaks were significantly associated with TE class DTM, while TE classes RLG and RLX showed significantly less peaks than expected (Figure 3A).

**Figure 2.**
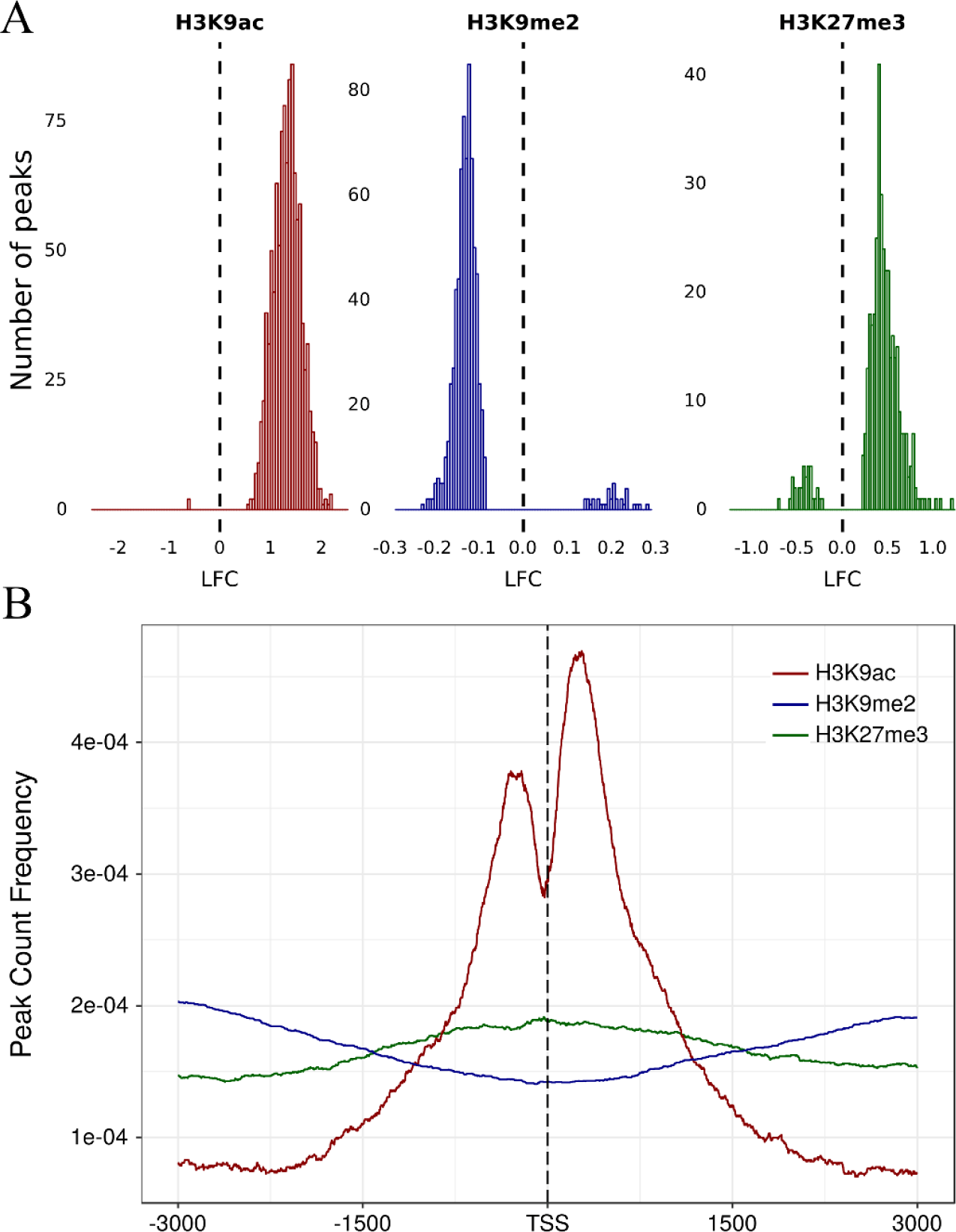
Analysis of differentially modified histone peaks in 3 dpi nematode-induced galls versus uninfected root tips. (**A**) Log2 fold change (LFC, infected vs. control) of significantly differentially enriched peaks for the 3 studied histone marks, (**B**) Peak distribution at the region 3 kb upstream to 3 kb downstream of transcription start sites (TSS) for the three studied histone marks.

**Figure 3.**
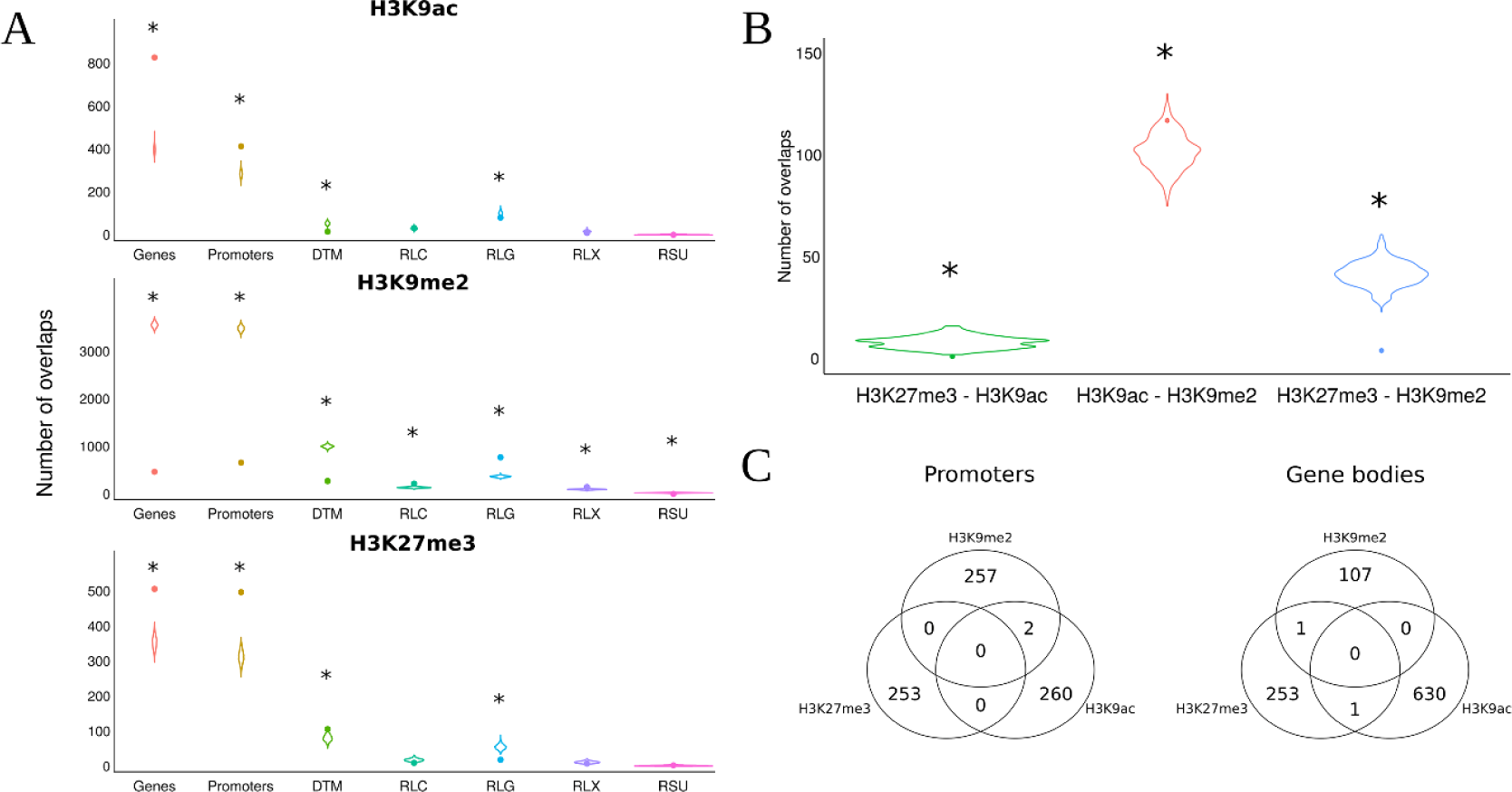
Concurrence of histone marks detected in 3 dpi nematode-induced galls and uninfected root tips. A. Significance of overlap between histone peaks and different genic regions or transposable elements (TEs). The violin plots show the distribution of the number of overlapping histone peaks and randomly scattered TE or gene regions (1000 simulations) and the dots the observed number. Asterisks indicate significant differences between the observed number and the random distribution (p < 0.05), (**B**) Similar as panel (**A**), but for the overlap between the different histone peaks, (**C**) Venn diagrams of the number of promoter regions and gene body regions shared between the three studied histone marks.

In total, 631 and 262 protein coding genes were hyperacetylated in their gene body or promoter, respectively (Supplementary file 3), significantly more than randomly expected (Figure 3A). When focusing on genes with H3K9ac peaks in their promoters no significant GO-terms were found and the WRS test (Benjamini Hochberg corrected) in MapMan also showed no enriched pathways. However, GO-analyses on genes with H3K9ac-peaks in their gene body revealed over-representation of genes related to protein modification, more specifically phosphorylation, response to organic substance and sequence-specific DNA-binding (Figure 4). MapMan confirmed significant enrichment in Bin 30.2, containing ‘signaling receptor kinases’ (number of genes: 23, P-value=0.02).

**Figure 4.**
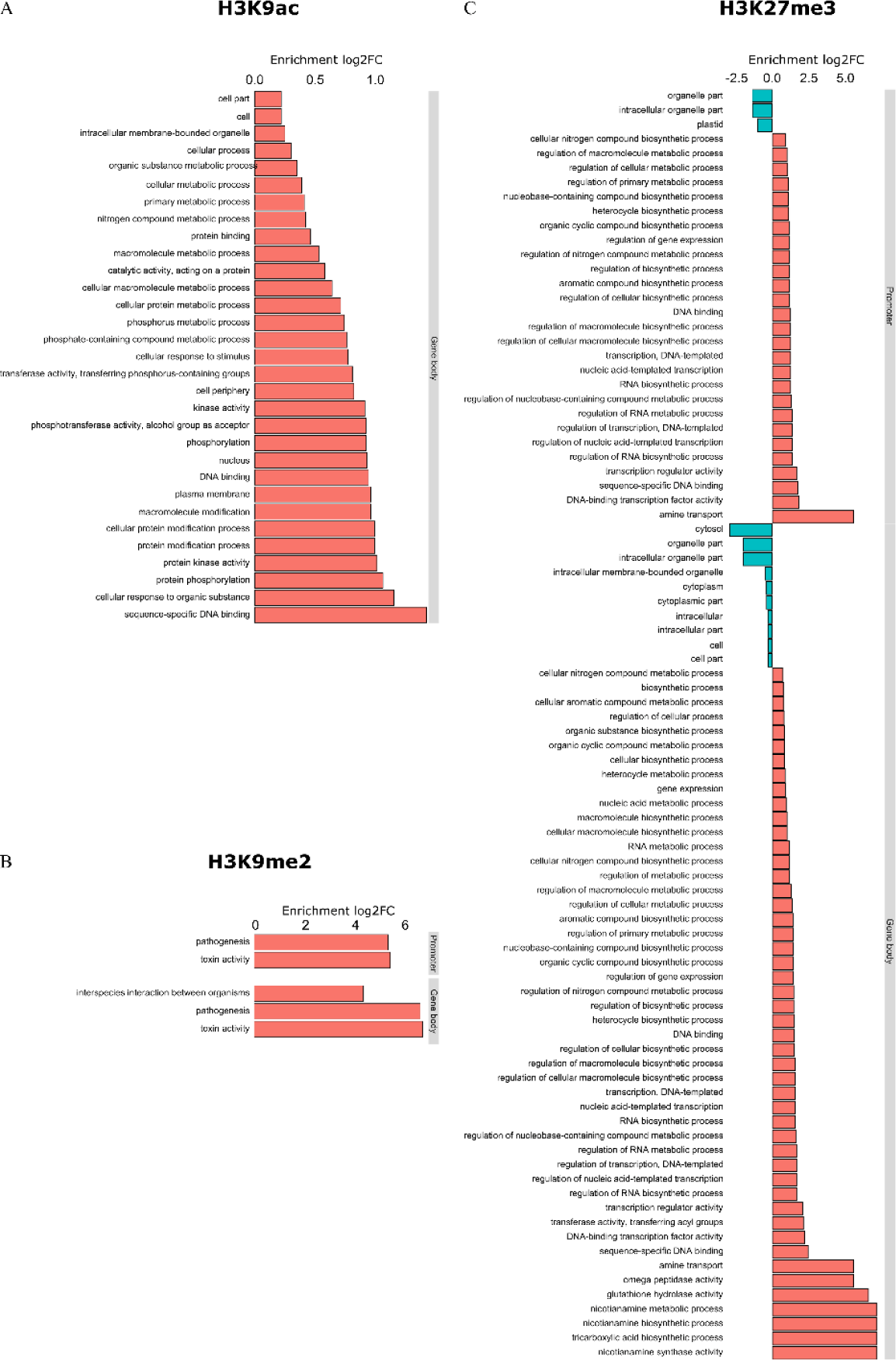
Gene ontology (GO) analysis of the genes associated with differentially modified histone peaks in 3 dpi nematode-induced galls versus uninfected root tips. (**A**) H3K9Ac (**B**) H3K9me2 and (**C**) H3K27me3. The graphs show the significantly enriched (Log2FC > 0) or depleted (Log2FC < 0) GO terms detected among the genes overlapping with the detected peaks in either gene body or gene promoter. Note that no significant GO terms were found in the set of genes that overlap with H3K9ac in their promoters.

Interestingly, a set of histone modifying genes like SET domain containing proteins and histone acetyltransferases were enriched for H3K9ac in their promoters or gene bodies. Many of the differentially modified peaks covered genes that are involved in plant defence, e.g. encoding PR-proteins, transcription factors of the WRKY, MYB, and ERF families, receptor kinases, MAP kinases and hormone related genes especially abscisic acid (ABA), auxin and ethylene (ET) (Figure 5 and Supplementary file 3).

**Figure 5.**
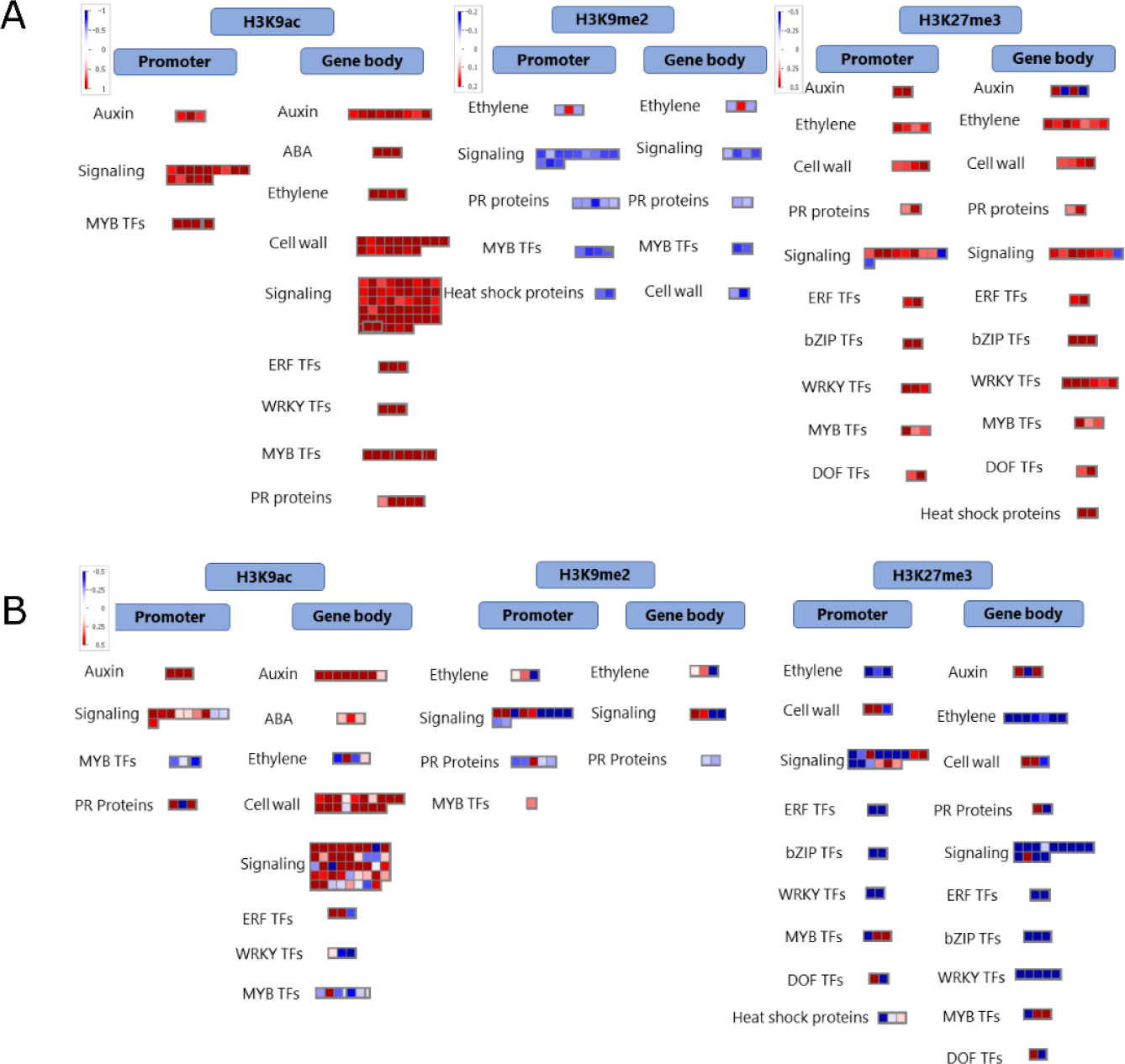
MapMan visualization of genes which are associated with significantly affected histone peaks in 3 dpi galls versus uninfected root tips in rice. (**A**) The visualization shows the pattern of the histone mark upon *Meloidogyne graminicola* infection in galls compared to uninfected control root tips. Red and blue denote enrichment and depletion respectively of this mark in infected roots *vs*. uninfected root tips. (**B**) Expression profile of these genes in 3 dpi galls *versus* uninfected root tips in rice (based on data published in Kyndt et al., 2012). Red and blue denote gene up- and down-regulation in galls *vs*. uninfected root tips, respectively.

### Nematode–induced galls feature depleted levels of H3K9me2

H3K9 methylation, including H3K9me2, is a repressive histone mark previously described to be involved in plant defence (Dutta et al., 2017). When comparing the presence of this mark in galls versus root tips we found a total of 732 genomic regions to be differentially methylated (Supplementary file 4). A majority of these peaks showed hypomethylation in the galls (Figure 2A). However, this decrease in methylation is less pronounced around TSS regions compared to H3K9ac (Figure 2B). Only 13 genes were found to have a hypermethylated promoter versus 246 genes with a hypomethylated promoter (Supplementary file 4). When focusing on the gene body, 8 genes were observed to be hypermethylated in H3K9 versus 100 genes hypomethylated (Supplementary file 6). Next to the significant depletion of this mark in gene bodies and promoters, this was also the case for TE classes DTM and RSU. On the other hand, positive associations were found between H3K9Me2 peaks and TE classes RLC, RLG and RLX (Figure 3A). The WRS test analyses in MapMan showed that there is no enrichment for any specific pathway among the genes with H3K9me2 in their promoter or gene bodies regions. However, H3K9me2 peak containing genes were strongly enriched for GO-terms related to pathogenesis and toxin activity (Figure 4). More specifically, we detected this mark to be associated with several genes involved in plant defence, such as cell wall synthesis, *PR* genes, thionin genes, transcription factors of the MYB family, receptor kinases, as well as hormone related genes for ethylene (Figure 5 and supplementary file 6).

### Nematode–induced galls have increased levels of H3K27me3

Galls and corresponding uninfected root tips were also used for ChIP-seq analyses targeting H3K27me3. Data analysis of the sequenced DNA revealed that upon nematode infection 496 peak regions were significantly differentially methylated (Supplementary file 5). The H3K27me3 peaks mainly show hypermethylation (Figure 2A) and their presence is increased around the TSS region (Figure 2B). Hypermethylation and hypomethylation of promoters occurred in 241 and 12 genes, respectively. Similarly, hypermethylation and hypomethylation of gene bodies occurred in 241 and 14 genes, respectively (Supplementary file 5). Next to the generally positive association between H3K27me3 and gene bodies as well as promoters, this was also the case for TE class DTM. A negative association was found for the presence of H3K27me3 peaks in TE classes RLC and RLG (Figure 3A).

The WRS test analyses in MapMan showed that no specific pathway is enriched among the genes with differential H3K27me3 peaks in promoters or gene bodies. However, GO-analyses revealed a large set of significant GO-terms related to transcription (factors), glutathione hydrolase and to nicotianamine biosynthesis (Figure 4). The list of genes associated with hypermethylated H3K27me3-patterns contains many well-known players in plant defence including transcription factors from the MYB, ERF and WRKY families, heat shock proteins, pectinesterase and thaumatin (Figure 5 and Supplementary file 5).

### Concurrence of studied histone marks

We evaluated whether the investigated histone marks concurred on genomic regions. Presence of H3K27me3 marks showed a significantly negative association with the H3K9ac-peaks and the H3K9me2-peaks. H3K9ac and H3K9me2 peaks featured a moderate but significantly positive overlap, while exhibiting opposite levels of modification changes between galls and roots i.e. an increase in H3K9 acetylation for a decrease in methylation and vice versa (Figure 2A, Figure 3B). When evaluating solely gene bodies or promoters (Figure 3C; Supplementary file 6), concurrence between marks was less clear, most likely due to the smaller part of the genome considered.

### Only minor correlation between differentially modified histones and genome-wide gene expression profiles in galls induced by *Mg*

H3K9ac has been typically associated with transcriptional activation while H3K9me2 and H3K27me3 are mainly associated with transcriptional repression. Here, we created a total RNA seq dataset to see whether there is a genome-wide correlation between differentially modified histones and gene expression profiles in *Mg*-induced galls at 3 dpi. Although some correlations were found to be statistically significant, the generally low correlation values and the appearance of the correlation plots make it unlikely that these correlations are biologically relevant (Figure 6). A significantly lower than randomly expected number of differentially expressed genes was found among the genes with differentially modified H3K9me peaks in promoter or gene body (Figure 7).

**Figure 6.**
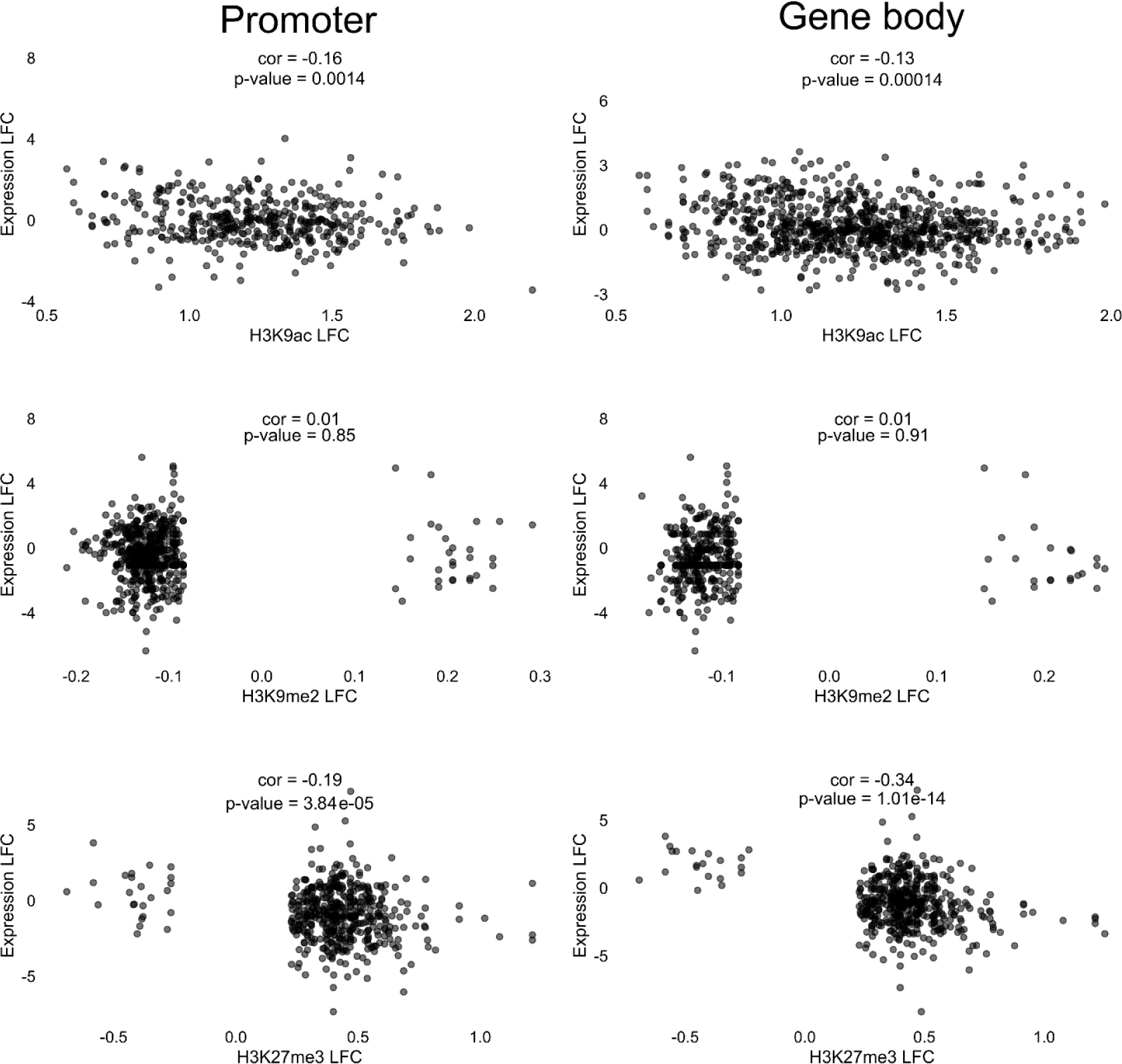
Correlation analyses between differentially modified genes and gene expression in nematode-induced galls. Correlation analyses between differentially modified H3K9ac, H3K9me2 and H3K27me3 peaks between 3 dpi galls and uninfected root tips, in comparison with the gene expression profile of the associated genes at the same time point. These analyses were performed separately for peaks detected to be associated with gene promoters (left) or with gene bodies (right).

**Figure 7.**
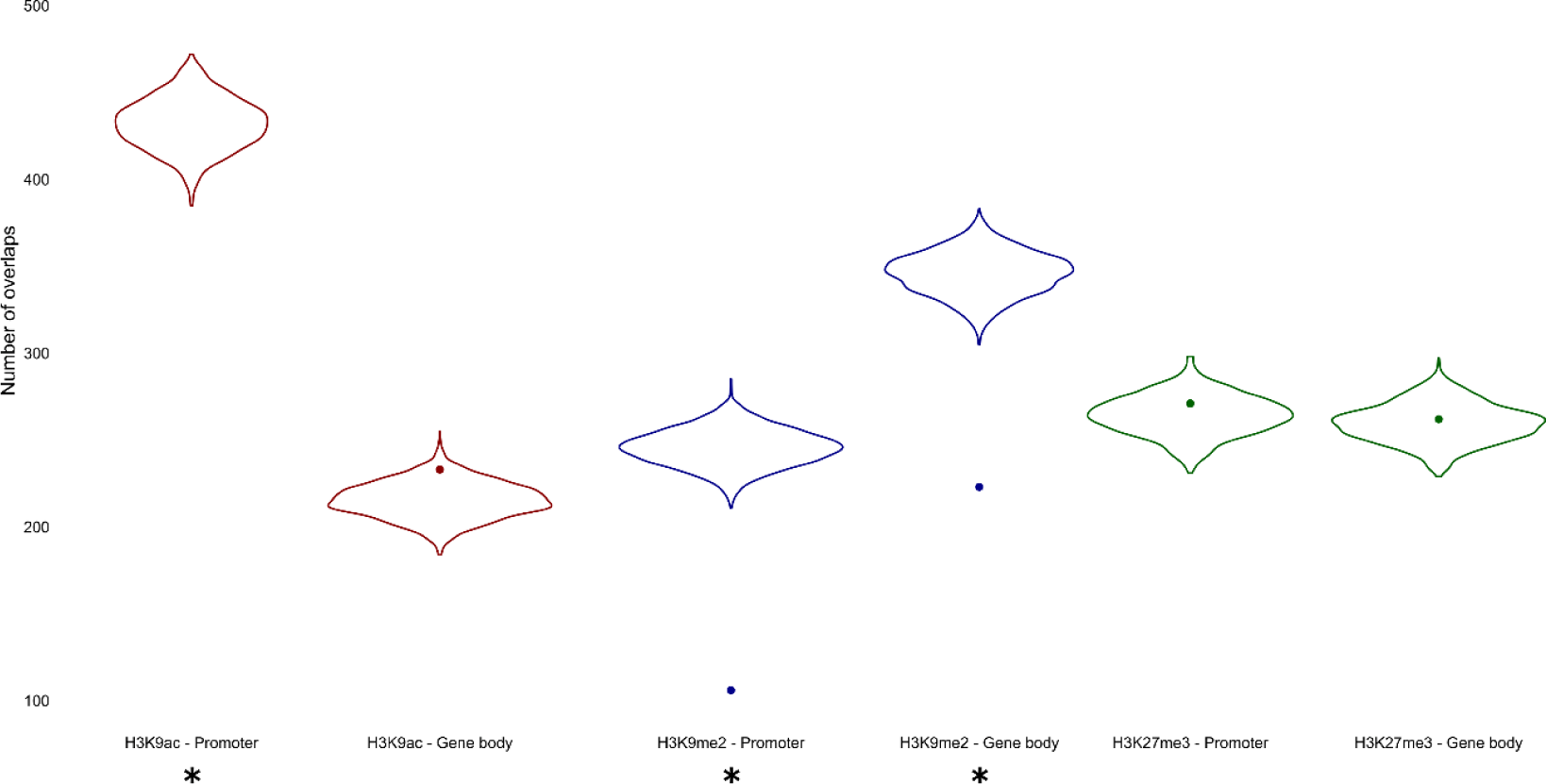
Association between differentially expressed genes and differentially modified histone peaks. The violin plots show the background distribution of overlaps between differentially modified peaks and a randomly sampled group of genes/promoters. The dots show the here-observed number of overlaps between differentially modified peaks and DE genes, promoters of DE genes. Asterisks indicate significant associations (P-value < 0.05).

Although an enrichment for differentially expressed genes was found among the genes with differentially modified H3K9ac peaks (Figure 6), no tendency towards upregulation was detected, as would be expected if the H3K9ac mark would be an activating histone mark. Of the 233 DE genes whose promoter overlaps with a differentially modified H3K9ac peak, 106 were upregulated (p = 0.93). DE genes with a differentially modified H3K9ac peak in their gene body do show a near significant trend towards upregulation: of the 543 DE genes whose gene body overlaps with a differentially modified H3K9ac peak, 291 were upregulated (p = 0.051).

Details about differentially expressed genes and their overlap with differentially modified peaks can be found in supplementary file 7.

### Association between histone marks and plant defence genes

Despite the genome-wide lack of clear correlation between histone peaks and differentially expressed genes, the GO-analyses indicated an enrichment for plant defence related genes. Fifteen genes were selected for qRT-PCR based validation of the expression patterns in independently collected *Mg*-induced gall samples at 3 dpi. For five defence-related genes associated with significant H3K9 hyperacetylation and gene activation, activation patterns were generally confirmed by qRT-PCR (Table 1A), validating the general robustness of our data and the correlation between this mark and defence gene activation.

**Table 1.**
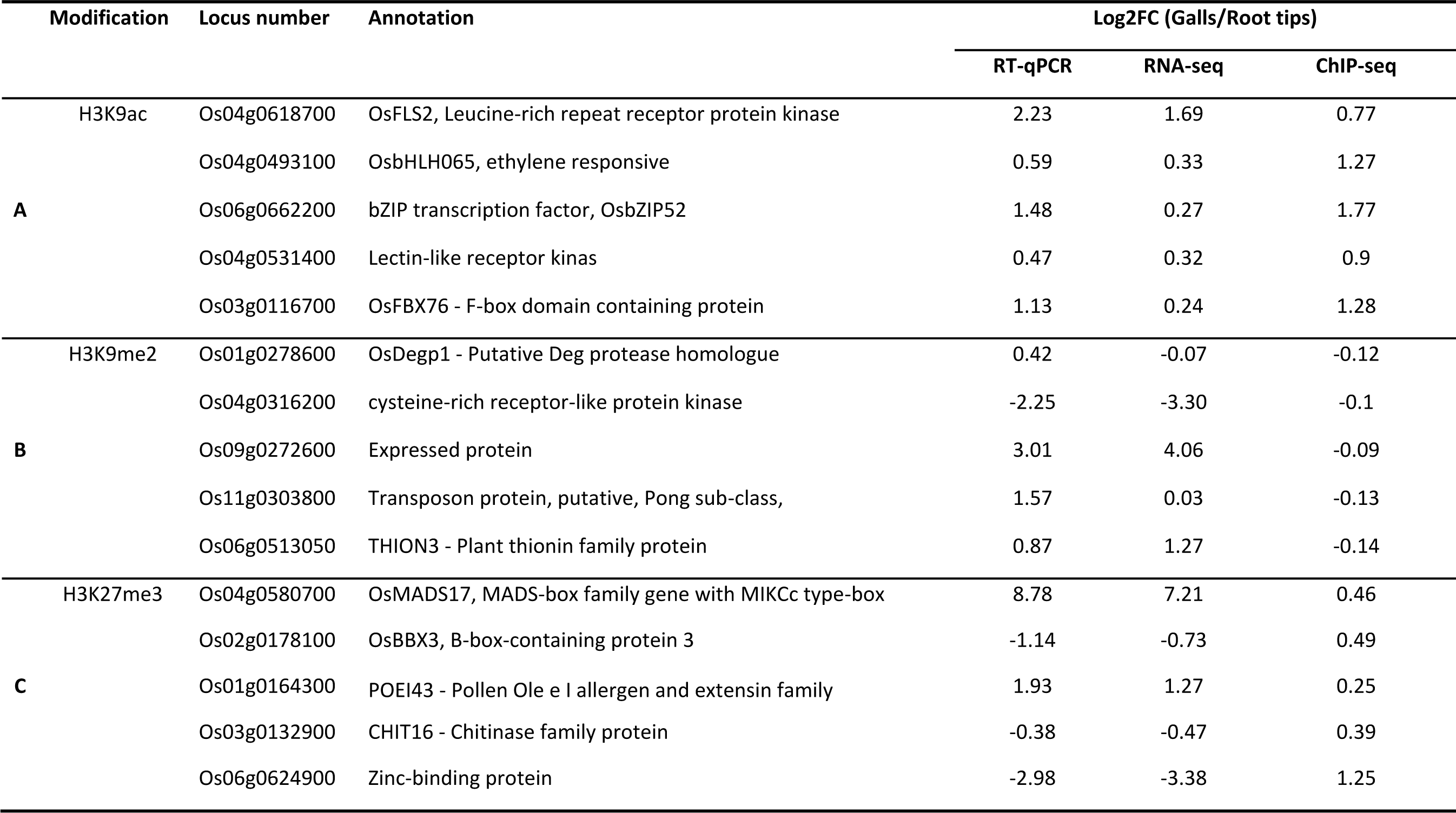
qRT-PCR based confirmation of the expression pattern of a subset of genes which are associated with differential histone marks as well as differential gene expression levels in 3 dpi galls induced by root-knot nematodes in rice in comparison with uninfected control root tips. The values indicate the mean of three biological replicates. Each biological replicate consists of a pool of galls obtained from about 20 plants.

For five defence-related genes that were significantly associated with H3K9me2 hypomethylation and that showed gene activation, qRT-PCR generally corresponded with the transcriptome results, except for 1 gene (Table 1B). This reveals that depletion of this mark may be slightly correlated with gene activation.

Five defence-related genes associated with significant H3K27 hypermethylation were selected. These genes showed diverging expression profiles in the RNA-seq data, and this pattern was here independently confirmed by qRT-PCR on gall samples (Table 1C). These data reveal no clear correlation between presence of this mark and gene expression.

## Discussion

Infection of rice plants by root-knot nematode *M. graminicola (Mg)* was previously reported to cause a striking modulation in the expression of genes involved in histone modifications (Kyndt et al., 2012; Ji et al., 2013; supplementary figure 1). Here, we studied the role of histone modifications in plant defence responses against *Mg*, and provide the first genome-wide data of histone modification patterns in the interaction between rice plants and these parasitic nematodes. At the investigated time point, 3 dpi, early giant cell formation is observable in rice roots and galls become macroscopically visible allowing specific sampling of the local infection site (Mantelin et al., 2017).

Our experiments with chemical inhibition and ectopic expression of histone modifications revealed contrasting results on plant susceptibility against *Mg* infection, suggesting a complex role of histone marks in controlling the plant defence response. Nevertheless, application of these chemicals could have widely affected the expression of many histone modifying enzymes and consequently defence and development. We assume that these chemicals generally affect the primary metabolic flux and pool of precursors for many development as well as defence-related - processes in plants, leading to reduced susceptibility for *Mg* in rice. For example, it has been shown that application of sulfamethazine reduces the plant folate pool and consequently the levels of S-adenosyl methionine, the donor of methyl groups for many substrates (Shen et al., 2016). Similarly, the lack of knowledge on the specific target regions of the enzymes over-expressed in the used transgenic lines precludes drawing specific conclusions about the reason for their affected susceptibility to nematodes.

To gain more specific insights into the changed profiles upon nematode infection, we studied the genome-wide pattern of three histone marks namely H3K9ac, H3K9me2 and H3K27me3, with the first one generally considered as a gene activation mark and the two latter gene repression marks (Armstrong, 2014). Confirming its role in gene activation the generally hyperacetylated H3K9ac peaks were mainly detected around the TSS site. The repressive mark H3K9me2 was mainly detected in non-genic regions, such as TEs, but was significantly under-represented in genes, promotors and around the TSS (Figure 2 and 3A). In case of H3K27me3, the generally hypermethylated peaks were mostly detected around the TSS. Only a small number of genes showed significant changes for more than one of the here-investigated histone marks (Supplementary file 6).

Our data revealed a strong genome-wide quasi-unilateral shift in these histone marks early upon *Mg* infection, with general enrichment of genome-wide H3K9ac, and H3K27me3 but depletion of H3K9me2 in young gall tissue compared to uninfected root tips. Strong genome-wide modification of histone marks in plants upon pathogen infection has been described before, but only upon aboveground infection by phytoplasma and fungi. For example, in bean plants infected with rust pathogen *Uromyces appendiculatus*, H3K9me2 and H4K12ac showed strong unilateral modification and similar to our data, transcription factors from the WRKY, MYB and bZIP families were affected by these marks (Ayyappan et al., 2015). In another study on *Paulownia fortune* upon infection by phytoplasma, 1,788 and 939 genomic regions were hyper- and hypoacetylated at H3K9 (Yan et al., 2019). Although a sharp hyperacetylation was not observed the peaks reported in that study were also related with GO terms metabolic process, cellular process and response to stimulus, as seen in our data (Figure 4) (Yan et al., 2019). As expected based on the results of the chemical inhibitors and transgenic lines, many defence related genes were marked by the studied histone marks (Figure 5). For example, many plant defence related genes were strongly hyperacetylated at H3K9, which is correlated with activated expression of these modified genes (Table 1A).

In contrast to H3K9me2, H3K9ac and H3K27me3 peaks were strongly associated with gene promoters and gene bodies (Figure 2). However, no clear genome-wide correlation with transcriptome data of 3 dpi galls was observed for any of the histone marks (Figure 6). This observation is confirming a previous study where no general correlation between loss or gain of H3K27me3 and gene expression was detected in the rice inflorescence meristem (Liu et al., 2015). We therefore hypothesize that there is a sophisticated cross talk between multiple epigenetic marks in the regulation of gene expression in plants. According to our observations there are strong modulations in the histone acetylome and methylome upon *Mg* infection, although this is not associated with genome-wide changes in expression profiles at the same time point. Further studies using other histone marks and multiple time points will be required to elucidate if and how the network of histone modifying genes, resulting marks and their interactions can modulate the genome-wide expression of the targeted genes. For example, it is possible that the genes are primed for activation at later time points.

Nevertheless, we have detected specific changes in histone marks around a subset of plant-defence related genes for each investigated mark. Also, the unilateral change in histone dynamics for each of the here-studied marks indicates that histone modifications play a role in the plant response upon *Mg* infection. The question arises how the genome is able to target histone changes specifically towards a subset of defence-related genes, and whether the parasitic nematode actively interferes with this response to attain susceptibility. We speculate that each histone modifying enzyme has the specific task to control a certain group of plant defence genes, such as for example shown for HDAC19 in controlling JA/ET-based defence responses (Zhou et al., 2005). Further detailed functional investigations on specific HDAC, HMTs and SDGs will be needed to prove this hypothesis.

## Conclusions

In summary, in this manuscript we present the first genome-wide study of histone marks upon *Mg* infection in plants. We showed that each histone mark showed a different unilateral shift (hypo- or hyper methylation or acetylation) and many of the associated genes are involved in important defence pathways. Chemical or genetic interference with histone modifying enzymes shows that these changes affect the susceptibility of plants towards infection, although with varying results, potentially depending on the diverging downstream defence pathways affected by this specific histone mark or modifying enzyme. Additional research using other histone marks and the elucidation of the exact underlying defence mechanism activated upon chemical or transgenic modulation of specific histone modification pathways are needed to depict a complete picture of epigenetic mechanisms upon infection of plants with *Mg* and potentially other pathogens.

## Materials and methods

### Plant growth conditions

*Oryza sativa* L. cv. ‘Nipponbare’ (GSOR-100, USDA) seeds were germinated for 5 days in darkness at 30 °C, after which they were transferred to synthetic absorbent polymer (SAP) substrate in polyvinylchloride tubes (Reversat et al., 1999) and grown at 28 °C (16 h light/8 h darkness). Plants were watered by Hoagland solution every other day.

### Isolation of activation-Tagged Mutants of Os*SDG740* and Os*HDA712*

Two activation tagging lines and their corresponding wild-type were obtained from the Postech PFG T-DNA insertion library, Department of Plant Systems Biotech, Kyung Hee University, Korea (Jeon et al., 2000; Jeong et al., 2006). These lines are activated in gene *OsSDG740* (PFG_K-03554.L; Os08g10470, cv. ‘Kitaake’) or in *OsHDA712* (PFG_2B-40507.L; Os05g36920, cv. ‘Dongjin’). T-DNA insertion and expression level of the target gene was confirmed using PCR and qRT-PCR, respectively (see further), using the primers given in Supplementary table 1. The *OsSDG740* and *OsHDA712* activation lines revealed a 16 and 11.5-fold induction of this gene in comparison to their corresponding wild-type, respectively.

### Vector construction and rice transformation for overexpression of *OsSDG729*

*OsSDG729* (Os01g56540) was amplified from DNA of *Oryza sativa* L. cv. ‘Kitaake’ using a high fidelity Taq DNA polymerase (Bioline, VELOCITY DNA Polymerase) and cloned into entry vector pDONR221 (Addgene) using Gateway BP clonase (Thermo Scientific). After confirmation by sequencing, the fragment was subcloned into binary vector PMBb7Fm21GW-UBIL (VIB, UGent, Belgium) using Gateway LR clonase (Thermo Scientific) and introduced into *Agrobacterium tumefaciens* EHA105 cells using tri-parental mating. Rice transformation was performed according to Langdale’s lab protocol adapted for Kitaake rice (https://langdalelab.com/protocols/transformation/). Transformed calli were selected on Luria Broth medium supplemented with 35 mg/mL of DL-Phosphinothricin (Duchefa, The Netherlands). T1 generation plants were selected by spraying 30mg/L of DL-Phosphinothricin. For evaluation of the expression level of *OsSDG729* in T1 transgenic plants, qRT-PCR was performed (see further). Two independent lines with the highest induction pattern, namely 128-fold (line F3-5) and 16-fold (line B1-6) induction in comparison with wild-type were selected. All primers are listed in Supplementary file 1.

### Nematode culture and inoculation

A pure culture of *M. graminicola* (*Mg*) was originally obtained from the Philippines (kindly provided by Prof. Dirk De Waele, Catholic University Leuven). The nematodes were cultured on susceptible rice and grasses (*Echinocloa crus-galli*). Nematodes were extracted from 3-4 month old cultures using the tray method (Whitehead and Hemming, 1965). After 2-3 days, the nematode suspension was collected and concentrated using 20 µm sieves. Two-week-old plants were inoculated with about 300 second-stage juveniles per plant or mock inoculated with water.

For infection assays, the level of infection was evaluated two weeks after inoculation. Root and shoot lengths and weights were measured. Root systems were harvested and stained by boiling in 0.013 % acid fuchsin for 5 minutes. After destaining in acid glycerol, the total number of galls and nematodes were counted under a stereomicroscope SMZ1500 (Nikon).

### Application of chemical inhibitors and evaluation of plant susceptibility

Thirteen-day-old rice plants (*Oryza sativa* cv. Nipponbare) were sprayed with nicotinamide, sulfamethazine or fumaric acid, while control plants were mock-sprayed with water. The surfactant Tween 20 was added to all spraying solutions at 0.02 % (v/v). Twenty four hours later, plants were inoculated and infection level was evaluated as described above.

### Data analysis

For statistical analyses software SPSS (Vesion25, IBM, USA) was used. The normality of data was checked using a Kolmogorov-Smirnov test (α=0.05). Homoscedasticity of data was verified using the Levene test (α=0.05). If data normality was confirmed, parametric tests, i.e. t-test, ANOVA and Duncan were used. Otherwise, non-parametric tests Mann-Whitney or Kruskal-Wallis, were applied.

### Synchronization of infection and chromatin immunoprecipitation (ChIP)

Two-week-old rice plants were grown and inoculated as described above. After 36 hours, when the majority of nematodes has entered the root system (Mantelin et al. 2017), they were transferred to 50 % Hoagland solution in glass tubes to synchronize the infection process. Three days post inoculation (3 dpi), galls of infected plants and uninfected root tips of mock-inoculated plants were harvested.

Chromatin extraction and shearing was performed according to the Diagenode Chromatin Shearing Optimization Kit for Universal Plant ChIP-seq Kit (Cat. No. C01020014) and precipitation according to the Plant ChIP-seq kit (Cat. No. C011010150) with some modifications. Briefly, collected galls and root tips (for each three biological replicates), were cross-linked in a buffer containing 1 % formaldehyde for 15 minutes on ice in a desiccator under vacuum condition. Glycine was added to stop crosslinking under an additional 5 minutes of vacuum. Then, root materials were ground and chromatin was extracted using three extraction buffers as in the manufacturer’s guidelines and collected and dissolved in 600 µL of sonication buffer supplemented with Protease Inhibitor Cocktail (Sigma). Sonication was performed using a Covaris M220 Focused-Ultrasonicator with following settings: Peak power of 75, duty factor of 10 and cycle/burst of 200 for 15 minutes. Sonicated plant materials were centrifuged (16,000 g for 5 minutes) and supernatant was collected. Chromatin immunoprecipitation was performed using three antibodies (Diagenode, Belgium), H3K9ac, H3K9me2 and H3K27me3. For chromatin precipitation, magnetic beads (DiaMag protein A-coated magnetic beads: Diagenode) were incubated with each antibody overnight, after which they were washed with ChIP dilution buffer to remove unbound antibodies. Beads were incubated for 10h with 50 µL of sheared chromatin diluted in ChIP dilution buffer (Diagenode, Belgium). One tenth of the diluted chromatin was collected and kept aside as an input sample before incubation with antibody. After incubation, beads were washed with washing buffers to remove unbound DNA fragments and DNA was eluted. Input and precipitated DNA were washed in absolute ethanol and the pellet was dissolved in 20 µL DNase free water.

### Library preparation and sequencing

Library preparation was done using the NEBNext Ultra II DNA Library Prep Kit for Illumina (New England Biolabs). To prepare samples, 34 µL DNase-free water was added to 16 µL precipitated DNA and adaptors were ligated to DNA fragments according to the manufacturer’s instructions (NEBNext Multiplex Oligos for Illumina kit NEB). After ligation, the DNA samples were cleaned up using AMPure XP beads (0.9x) (Beckman Coulter). Resulting fragments were amplified for 14 cycles using NEBNext Ultra II PCR protocol (New England BioLabs) and quality was assessed with the Agilent High sensitivity DNA kit. Amplicons were excised from a 2 % agarose gel (200-800 bp fragments were excised) and purified using the Gel DNA recovery kit (Zymo Research). Library concentrations were measured with qPCR according to Illumina’s Sequencing Library qPCR Quantification Guide. Sequencing was done on a NextSeq500 using single reads (76 bp). A 2.3 pM library was loaded on the flow cell with 2 % PhiX spike-in.

### Data analysis of ChIP-sequencing data

Reads were trimmed using Trim Galore (version 0.4.0) with default parameter settings. Mapping was done with Bowtie (version 1.1.1). Unmapped reads were filtered out and multiplexed subsamples were merged with samtools (version 1.9). Redundant reads were filtered out using Picard (version 2.18.27). For peak calling, the MUSIC (multiscale enrichment calling) software (Harmanci et al., 2014) was used. A mappability profile was generated using the *Oryza sativa* ssp. *japonica* (build MSU7.0) genome with a read length parameter of 76. For peak calling, all samples were merged per histone modification (excluding the input sample which was used as a control for the genome background). For histone modifications H3K9me2 and H3K27me3 the broad peak subroutine was used while for histone modification H3K9ac punctate peaks were called, following the classification of the ENCODE project (https://www.encodeproject.org/chip-seq/histone/).

Read quality was checked using FastQC (version 0.11.8). Sample quality was evaluated by calculating the non-redundant fraction and the PCR bottleneck coefficient. Successful enrichment of genomic regions was checked using the R package ChipQC version 1.18.2 (Carroll et al., 2014) and calculating the fraction of reads in peaks as well as the standardized standard deviation. The normalized strand cross-correlation coefficient and the relative strand cross-correlation coefficient were also calculated using phantompeakqualtools (Kharchenko et al., 2008; Landt et al., 2012). Read counts and quality control metrics can be found in Supplementary file 2. For histone modifications H3K27me3 and H3K9me2 three biological replicates were used per condition, for H3K9ac two biological replicates were used, because a third replicate was of insufficient quality.

To assess significant changes in peak regions between galls and roots, count tables of read counts in peak regions were generated by the summarizeOverlaps function in the GenomicAlignments R package (version 1.16.0). Differential peaks were identified using the DESeq2 package (version 1.22.2) with an FDR cutoff of 0.05. P-values were corrected for multiple testing using Benjamini-Hochberg correction (Love et al., 2014). Peak annotation of differential peaks and transcription start site analysis was done using ChIPseeker (version 1.18.0) (Yu et al., 2015). Gene and TE annotations were obtained from the Ensembl (Oryza_sativa.IRGSP-1.0.42.gff3) and Rice Transposable Element databases. Five TE classes were used: DNA Transposon Mutator (DTM), long tandem repeat Copia (RLC), long tandem repeat Gypsi (RLG), long tandem repeat unknown (RLX) and short interspersed nuclear elements (RSU). Promoter regions were defined as regions 2000 bp upstream of the transcription start site.

### Association between gene sets

Significance of association between sets of genomic regions was evaluated using permutation tests using the regioneR package (version 1.14.0) (Gel et al., 2016; Catoni et al., 2017; Zhao et al., 2019). First, 1000 permutations were performed to create a null-distribution. Per permutation, a set of genes of equal size to the set of genomic regions was randomly selected from the total set of rice genes and the number of overlaps was counted. Subsequently, the number of true overlaps between the two gene sets was compared to this null distribution (cut-off: p < 0.05).

### Gene expression profiles

Rice was grown and inoculated as described above. For each treatment (3 dpi galls or root tips), 3 independent biological replicates were sampled. RNA was extracted using the ZR Plant RNA Miniprep kit (Zymo Research) and DNAse-treated (ThermoFisher). RNA quality was verified with a RNA 6000 nanochip (Agilent technologies) and concentration was measured with a Quant-it Ribogreen RNA assay (Life technologies). For rRNA depletion, 1500 ng of RNA was treated with the Ribo Zero Plant Seed/Root kit (Illumina). The Truseq stranded total RNA library prep (Illumina) was used for library prep. The cDNA was used for enrichment PCR (13 cycles), purified with the double AMPure XP cleanup (1:1) (Beckman Coulter), and checked with a High sensitivity DNA chip (Agilent technologies). Quantification of the libraries was done with a qPCR assay according to Illumina’s protocol to enable equimolar pooling of libraries. Finally, sequencing was performed on a NextSeq 500 using 2% Phix spike-in (single end reads, 76 bp). Read quality was verified with FastQC (version 0.11.8), and reads were trimmed using Trimmomatic (version 0.38) with following parameters: ILLUMINACLIP:3:30:10, MAXINFO:23:1, SLIDINGWINDOW:5:30, MINLEN:17. STAR (version 2.6.1d) was used for mapping on the *Oryza sativa* ssp. *japonica* (build MSU7.0) genome with following parameters: readFilesCommand zcat, outFilterMultimapNmax 1 and outSAMtype BAM SortedByCoordinate. Afterwards, samtools (version 1.10) was used to merge multiplexed samples. A count table was made using rtracklayer (version 1.44.4) to convert the GTF (Ensembl release 42) file into a Granges object, Rsamtools (version 2.0.3) to create a BamFileList object and GenomicAlignments (version 1.20.1) for the summarizeOverlaps function to create the count table. To find differentially expressed genes, DESeq2 (version 1.24.0) was used with an FDR cutoff of 0.05. P-values were corrected for multiple testing using the Benjamini-Hochberg correction.

Correlation between changes in histone modifications and gene expression was evaluated by comparing Log_2_ fold changes of gene expression level between galls and control samples with the Log_2_ fold changes of their overlapping histone peaks using the Pearson correlation coefficient. Association between differentially expressed genes and differentially modified histone peaks – either associated with gene bodies or with promoters - was tested with the regioneR package as described above.

### Gene ontology and pathway enrichment

For gene ontology analyses, the PLAZA 4.5 platform with default parameters was used (with Bonferroni correction; p < 0.05,) (Van Bel et al., 2018). The Wilcoxon Rank Sum (WRS)-test (with Benjamini Hochberg correction, p < 0.05) was used to analyze the statistical enrichment for pathways in MapMan (Thimm et al., 2004).

### RNA extraction, reverse transcription and qRT-PCR analysis

RNA was extracted using the RNeasy Plant Mini Kit (Qiagen) and cDNA was synthesized using Tetro Reverse Transcriptase (Bioline). qRT-PCRs were performed with three technical and three biological replicates. The qPCR conditions consisted of initial denaturation at 95 °C for 10 min, followed by 50 cycles of [95 °C for 25 s, 58 °C for 25 s, 72 °C for 20 s]. Expression data were normalized using data of two reference genes (all primers are listed in Supplementary file 1) and analyzed using REST2009. Transcript levels in gall samples are expressed relative to the expression level in uninfected control root tips.

### Accession numbers

Data were deposited in the Gene Expression Omnibus: GSE145501 (Chip-Seq) and GSE152783 (RNA-seq).

## Supplementary data

**Supplementary figure 1.** Expression profile of genes involved in histone acetylation, deacetylation and lysine methylation and demethylation in nematode-induced galls at 3 dpi, as observed in our previously published transcriptome analysis of the rice-*Meloidogyne graminicola* interaction (Kyndt et al., 2012). The gene annotation is based on (Zhou et al., 2013). Genes selected for over-expression are indicated with an arrow.

**Supplementary figure 2.** Root and shoot length in overexpression lines of (A) *OsSDG729*, (B) *OsSDG740* and (C) *OsHDA712*. Data was taken at the end of the infection experiment shown in Figure 1J-L. Error bars indicate SEM.

**Supplementary file 1.** Primers used in this study.

**Supplementary file 2**. Read counts per sample and quality control metrics.

**Supplementary file 3.** List of genes associated with differentially modified H3K9ac peaks in their promoter or gene bodies at 3 dpi upon nematode infection in rice roots.

**Supplementary file 4.** List of genes associated with differentially modified H3K9me2 peaks in their promoter or gene bodies at 3 dpi upon nematode infection in rice roots.

**Supplementary file 5.** List of genes associated with differentially modified H3K27me3 peaks in their promoter or gene bodies at 3 dpi upon nematode infection in rice roots.

**Supplementary file 6.** List of genes associated with multiple detected histone marks in their promoter or gene bodies at 3 dpi upon nematode infection in rice roots.

**Supplementary file 7**. Differentially expressed genes after *Meloidogyne graminicola* infection 3 days post inoculation and their overlap with differentially modified histones.

## Responsibilities of the Author for Contact

The authors will ensure communication and share data upon request.

